# Diet predicts milk composition in primates

**DOI:** 10.1101/197004

**Authors:** Gregory E. Blomquist, Katie Hinde, Lauren A. Milligan Newmark

**Affiliations:** Department of Anthropology, University of Missouri; School of Human Evolution and Social Change, Arizona State University; Center for Evolution and Medicine, Arizona State University; Brain, Mind, and Behavior Unit, California National Primate Research Center; Associate Faculty, Anthropology, Mira Costa College

**Keywords:** lactation, nutrition, parental care, foraging ecology, energetics, allometry

## Abstract

Mammalian mothers pay heavy energetic costs to fuel the growth of their offspring. These costs are highest during lactation. Energy transmitted to offspring in the form of milk must ultimately come from the maternal diet, but there have been few comparative studies of the relationship between milk properties and mammalian diets. We used interspecific data on primate milk composition and wild diets to establish that concentrations of milk protein and sugar are predicted by diet independent of maternal mass, litter mass, and infant parking behavior such that increasing folivory or faunivory increases protein concentration but decreases sugar concentration. Milk energy density is unrelated to diet, though infant parking species do produce more energy-dense milk. While parking effects have been previously explained as a result of mother-infant separation, the mechanisms causing the relationship between nutrient packaging in milk and maternal diet are currently unclear. However, they likely reflect evolved differences in maternal energetics related to maternal foraging ecology, infant growth patterns, or the relative dietary abundance of nutrients costly to synthesize in milk among primate species.

## Introduction

Lactation is the most costly phase of parental care for female mammals (Dufour & Sauther, 2002; Gittleman & Thompson, 1988). High costs imply great opportunity for selection to mold interspecific differences to local ecology. However, quantitative studies of the ecological correlates of lactation biology are extremely rare, partly as a result of a justifiable focus on the influence of maternal-offspring interactions and body size on milk properties (Ben Shaul, 1962; Blurton Jones, 1972).

The importance of allometric scaling in milk yield and composition is well established (Riek, 2008, 2011; Lee, 1999; Martin, 1996, 1984; Oftedal, 1985, 1984). From empirical scaling relationships between maternal mass and milk yield (g) and between maternal mass and milk energy content (kcal), Martin (1984) showed the expected scaling of maternal mass and milk gross energy (kcal/g) or other composition variables (e.g. percent mass of protein) should be weakly negative. This is because the exponent for mass-yield scaling is greater than the mass-energy content exponent. A similar negative relationship is predicted based on scaling of neonate mass. Using a very limited database of mammals, Martin (1984) confirmed these negative scaling relationships in primates and ungulates but not carnivores — possibly because of the high availability of fat and protein in their animal diet. As this result suggests, a variety of other factors may affect milk properties beyond allometric scaling (Power et al., 2002).

Martin (1984) drew attention to the influence of altriciality versus precociality of mammalian neonates. Altricial neonates are provisioned with dense milk having high concentrations of fat and protein but little sugar, while precocial neonates are reared on more dilute milk with high sugar and less fat or protein (Jenness & Sloan, 1970). In general, primates fit this trend, producing dilute milk for their relatively precocial infants. However, there are exceptions or potential problems with placing milk properties entirely within the altricial-precocial dichotomy. Marine mammals (pinnipeds and cetaceans) produce precocial neonates but have milk with very high fat and protein concentrations and almost no sugar. A further exception is the similarity in milk between rabbits (which have altricial young) and hares (with precocial young) (Oftedal & Iverson, 1995). An additional complication to placing milk properties within the altricial-precocial dichotomy is its interaction with litter size. Most precocial neonates are singletons while altricial neonates tend to be members of larger litters. Litter size itself explains further variation, particularly in milk energy content (kcal) (Oftedal, 1984, 1985).

Milk composition relationships with litter size and altriciality/preociality also cause it to be influenced by relative size of neonates (Derrickson, 1992; Martin, 1984). Across mammals, neonate mass scales negatively allometrically with increasing maternal mass (Smith & Leigh, 1998; Harvey & Clutton-Brock, 1985). In primates, this relationship holds when higher order births are accounted for by calculation of litter mass. Small-bodied primates have relatively larger litter mass, though there is also a clear increase in relative litter mass from strepsirhines to haplorhines (Martin, 1984; Leutenegger, 1979). Some of this variation may relate to maternal strategies with relatively heavier litters reflecting greater prenatal investment. Postnatal investment could then be negatively, positively, or unrelated to relative litter mass depending on the duration and intensity of nutrient transfer during lactation (Lee, 1999).

Finally, suckling rates vary widely among taxa with predictable outcomes on milk composition (Ben Shaul, 1962; Blurton Jones, 1972). Species in which mothers are separated from infants for long periods of time tend to have milk with less water and higher fat and protein concentrations than species where mothers and infants are rarely separated and feeding is “on demand.” Indeed, Tilden et al. (1997) have demonstrated this effect of suckling rates in strepsirhine primate milk. For example, infant carrying *Eulemur* species produce dilute milk but lorises or galagos, which “park” infants for long periods, have much higher fat and protein concentrations.

Additional ecological correlates of milk properties (e.g. aridity, temperature, locomotion, fasting or hibernation, mating system, diet) have not been explored quantitatively in primates but have been suggested for other mammalian orders. For example, among carnivores relatively slow infant/litter growth rates in foli/herbivorous species (e.g. black bears, giant pandas) have been related to low milk energy content. This may directly reflect lower quality diet (pandas), or be confounded with maternal strategies to conserve resources when emerging from hibernation (black bears). Foraging ecology, rather than nutrient composition of the diet, may also influence carnivore milk. Despite an exclusively meat diet, cheetahs and lions have low predicted milk energy output. This is thought to be a maternal bet-hedging strategy against low success in capturing prey and/or perhaps long travel distances (Oftedal & Gittleman, 1989; Gittleman & Oftedal, 1987). Similar dietary correlates of milk composition have been noted in bats (Kunz & Hood, 2000; Kunz & Stern, 1995). Insectivorous species generally have higher fat and protein concentrations than frugivorous species, though allometric effects may confound this comparison.

Here, we emphasize diet with the general hypothesis that maternal nutritional or foraging ecology should be reflected in primate milk properties. Milk yield data are not available on enough primate species for quantitative treatment so we restrict our attention to proximate composition. We recognize that primates produce relatively dilute milk, delivered to few infants over a protracted infancy compared to other mammals (Oftedal & Iverson, 1995), but consider them to be an ideal taxon for their range of body sizes, locomotor habits, social systems, habitats, and diets. Relatively large infants with costly large brains make primate infants very costly, but their relatively low growth rates and long female reproductive careers may also allow for variation among primate species in how mothers invest in infants that are not possible in other mammalian groups (Pereira & Leigh, 2003). From this perspective, Leigh (1994) hypothesized that folivorous primates should have higher milk protein because they grow faster than other primates of comparable body size. He noted the high protein content of structural plant parts could enable faster growth (Lambert, 1998; Tanner, 1990) and that rapid attainment of adult size and early maturation might reflect reduced feeding competition in folivores (Janson & van Schaik, 1993). Alternatively, rapid growth might be facilitated by higher milk gross energy (kcal/g), which would be reflected primarily in milk fat concentration. Crudely, this would imply a more faunivorous diet enables more rapid growth. A null alternative is that primate mothers produce essentially the same kind of milk regardless of diet — leaving interspecific milk composition differences to be related to allometry, mother-infant contact, neonate or litter size, and other unknown factors.

## Materials and Methods

### Milk Composition

We collected data from the literature on milk composition and a variety of potential predictors. Milk composition data came from original references in Hinde & Milligan (2011). The few other reports on primate milk composition were not included because infant age was unknown or details of milk collection or analysis were unspecified or atypical (*Saguinus oedipus*, *Cercocebus* sp., and *Macaca radiata*, Jenness & Sloan, 1970; Laudenslager et al., 2010). We note the taxonomic and ecological breadth of the sample is limited with no data on pithicines, colobines, tarsiers, or many strepsirhine clades (Figure 1). With a few exceptions, milk composition is only available for captive individuals (Hinde & Milligan, 2011).

**Figure 1.**
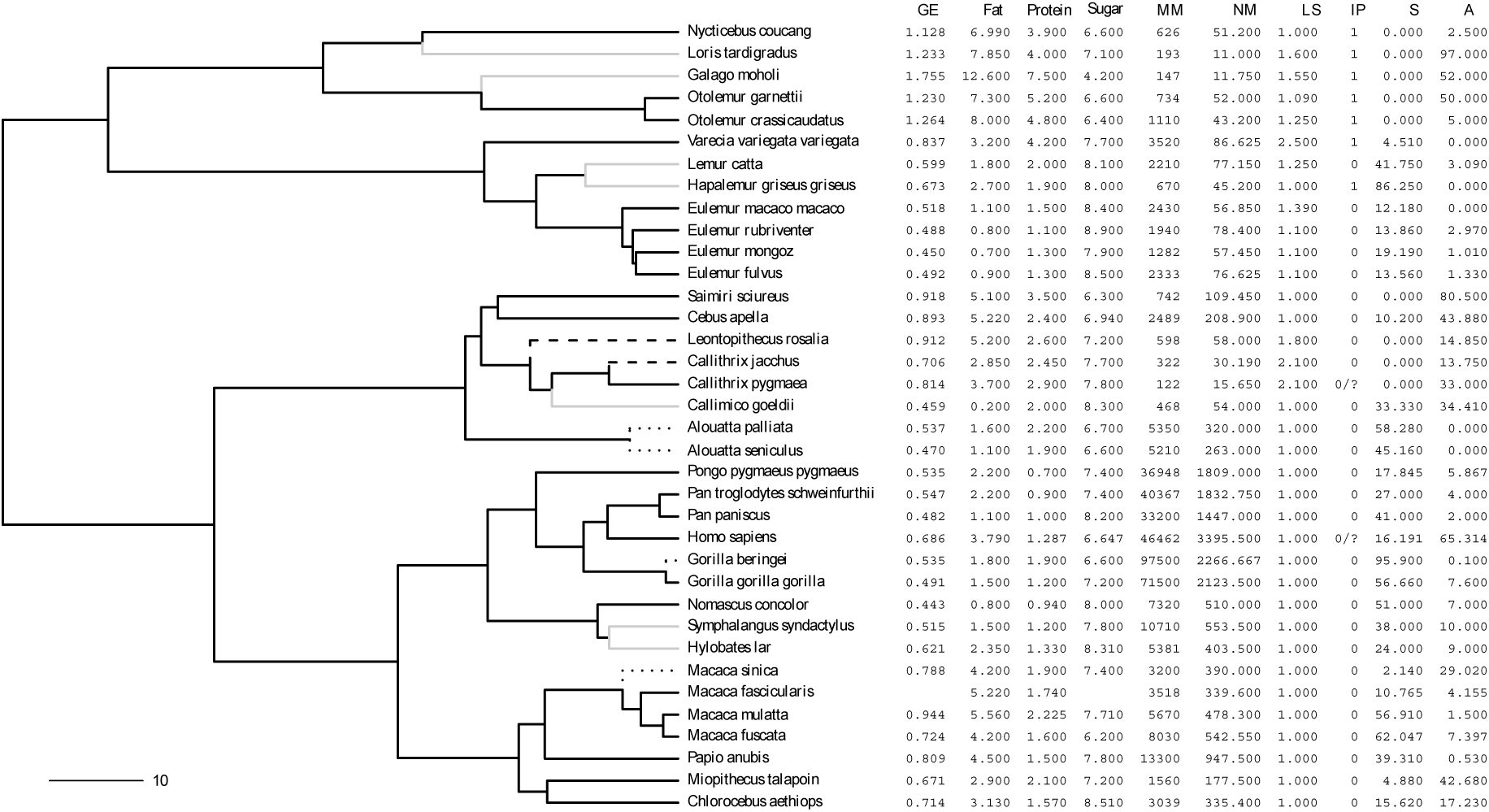
Primate phylogeny from 10,000 trees and Thalmann et al. (2007). Scale bar indicates 10 million years. Grey branches are species with a single milk sample. Dotted branches indicate wild-only milk data; dashed branches are for an average of wild and captive milk data. Values are given in the table: GE (kcal/g), percents fat, protein, and sugar (g/100 g), maternal and neonate mass (g), mean litter size, IP = infant parking (1 = parkers, ? = ambiguous), S = percent structural plant parts in diet, and A = percent animal matter in diet. A nexus file and spreadsheet of the tree and data are provided as supplements.

### Maternal mass and litter mass

Adult body mass was taken from Isler et al. (2008) using female values when available. *Nomascus concolor* is not included in their database so we used their congener *Nomascus gabriellae*. Adult human female mass was averaged from 29 cultures reported in Walker & Hamilton (2008). We used Smith & Leigh’s (1998) data on neonate mass, calculating an unweighted mean of sex and study for each species. Four species lacked neonate mass data and were supplemented by other sources. *Alouatta palliata* neonate mass was taken from Kappeler & Pereira (2003). For *Nomascus concolor* we used the “*concolor*”-group neonate mass value reported in Geissmann & Orgeldinger (1995). We used neonate mass for *Macaca sinica* of 390 g reported on ADW. Finally, we use new data on neonate mass for mountain gorillas (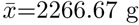, *N* = 3; C Whittier, personal communication). Mean litter sizes were collected from several sources (Roberts, 1994; Geissmann, 1990; Izard & Nash, 1988; Rasmussen, 1985; Harvey & Clutton-Brock, 1985), allowing litter mass to be calculated as the product of neonate mass and mean litter size.

### Diet

A species’ diet was quantified as percent time foraging on particular food items in the wild. Data were mostly found in the primary sources referenced in Campbell et al. (2011). Human diets are from foragers tabulated by Kaplan et al. (2000). The original categories were recorded and then summed into reproductive plant parts, structural plant parts, and animal prey. Percentages were rescaled to add up to 100%, if necessary. In the analysis we used percent structural and percent animal because they are likely to discriminate dietary habits in primates the most and, because the three percentages sum to 100, structural and animal percentages can be used together in multiple regression to give complete dietary information. Animal matter is probably the most important of these categories as a rich source of protein and fat. Structural parts are typically considered high in protein, and there is debate about variation in nutrient content of fruits, nectar, exudates, and flowers which are often considered high sugar, low fat and protein resources (Lambert, 1998). Seeds were included in reproductive parts even though they are high in fat and protein. Seed ingestion often cannot be discriminated from fruit flesh or flower consumption in the field. A data spreadsheet and full set of original references are provided as supplementary material.

### Maternal Care Patterns

Currently, there are few systematically collected data on species-typical patterns of nursing frequency and duration which prevents more nuanced investigation of maternal care and milk composition (Tilden & Oftedal, 1997). We therefore used infant parking as a proxy measure. Parking categories were coded from descriptions collected in Ross (2001) and Kappeler (1998). We used a binary code for frequency of broken contact between mother and infant based on parking of infants inside and outside of nests. This approach is crude but should capture any large differences between parking and non-parking species. While it would be better to have quantitative measure of duration of broken contact that could be matched directly to the infant age when milk were collected, this is not available for nearly all species. Despite reports of infant parking in marmosets (especially *Callithrix pygmaea*) we coded then as non-parking, considering this a recently derived variation on the callitrichine pattern of non-maternal infant carrying such that their milk would be unlikely to reflect parking behavior considering the strong phylogenetic signal in milk composition (see discussion of *λ* below) (Heymann & Soini, 1999; Digby & Barreto, 1996; Garber, 1997). Similarly, we coded humans as non-parking, interpreting this as the ancestral great ape pattern — though mother-infant separation is common in many cultures (Konner, 2005; Sellen & Smay, 2001). Because of the ambiguity of how to code these species, we also report a separate analysis excluding them in supplementary material.

### Data Analysis

We analyzed the relationship between milk composition and predictors by multiple regression of the species values (TIPS) and phylogenetic generalized least squares (PGLS) (Martins & Hansen, 1997; Grafen, 1989). This allows simultaneously testing the effects of multiple predictors and controlling for correlations among them (Freckleton, 2002). Milk composition, maternal mass, and litter mass were log_10_-transformed. Infant parking was entered as a dummy variable such that its slope estimates are the difference between parking and non-parking species. The PGLS models adjust for non-independence among the individual species arising from shared evolutionary history through a correlation matrix derived from an independently estimated phylogeny. Phylogenetic relationships among the primate species were taken from a consesus tree of version 2 of the *10,000 trees project* based on 1000 MCMC samples (http://10ktrees.fas.harvard.edu/). *Gorilla beringei* was added to the phylogeny with a divergence from western lowland *G. gorilla* at 1.2 million years ago (Thalmann et al., 2007). All analysis was conducted in R (R Development Core Team, 2009). TIPS models were run with the lm() function while the PGLS models were implemented in by gls() from the nlme package (Pinheiro et al., 2009). In the PGLS analysis, λ was estimated from the data in each model (Freckleton et al., 2002). This measure of the importance of phylogeny, ranging from 0 to 1, was high in all cases (≥0.9) indicating a strong phylogenetic signal. In TIPS and PGLS analysis, the combined effect of structural plant and animal percentages in the diet was tested by comparing a model including these two diet variables and a model excluding them with an *F*-test implemented in the base R function anova().

Because the milk composition data vary dramatically in the number of individual samples that contributed to the average values reported, we examined the full data set and a reduced data set that excluded several species that had data from a single milk sample only (Table 1). In addition, several outlier points were excluded from analysis of the full data set based on milk composition – adult mass plots (Figure 2). *Hapalemur* was excluded for all four milk variables because it was a major outlier of the infant parking group. *Callimico* and *Galago moholi* were excluded for fat and sugar, respectively, for which each has very low values. All three of these taxa were also excluded from the reduced data set because their milk data are from a single sample. Knowing that infant parking has strong effects on milk composition (Tilden & Oftedal, 1997), we ran separate analyses on the non-parking species alone but report the results in supplementary material because patterns were identical to the parker/non-parker combined analysis. Quantitative treatment of the parking taxa alone was not feasible with so few species.

**Table 1.** Multiple regression (TIPS) and phylogenetic generalized least squares (PGLS) models for primate milk composition.

|  | full data set |  | reduced, milk samples > 1 |  |
| --- | --- | --- | --- | --- |
| | TIPS<br>$\beta$ (SE) | PGLS<br>$\beta$ (SE) | TIPS<br>$\beta$ (SE) | PGLS<br>$\beta$ (SE) |
| GE (kcal/g) | full $N=33$, tips $R^2=0.802$, tips diet $F$ $P=0.26$, gls diet $F$ $P=0.47$<br>reduced $N=28$, tips $R^2=0.741$, tips diet $F$ $P=0.313$, gls diet $F$ $P=0.632$ | | | |
| (Intercept) | 0.1034 (0.0949) | 0.0100 (0.1383) | 0.0689 (0.1009) | -0.0956 (0.1569) |
| log10(MM) | -0.3099 (0.0749) *** | -0.2171 (0.1137) | -0.2905 (0.0813) ** | -0.1696 (0.1283) |
| log10(LM) | 0.3157 (0.0868) ** | 0.2329 (0.1439) | 0.2990 (0.0980) ** | 0.2043 (0.1655) |
| IP | 0.2810 (0.0429) *** | 0.2230 (0.0676) ** | 0.2796 (0.0489) *** | 0.2322 (0.0774) ** |
| S | 0.0008 (0.0008) | 0.0004 (0.0008) | 0.0008 (0.0009) | 0.0001 (0.0008) |
| A | 0.0010 (0.0006) | 0.0008 (0.0006) | 0.0013 (0.0009) | 0.0008 (0.0008) |
| Fat (g/100 g) | full $N=34$, tips $R^2=0.708$, tips diet $F$ $P=0.398$, gls diet $F$ $P=0.481$<br>reduced $N=29$, tips $R^2=0.653$, tips diet $F$ $P=0.58$, gls diet $F$ $P=0.547$ | | | |
| (Intercept) | 0.8805 (0.2479) ** | 0.5088 (0.3588) | 0.8204 (0.2719) ** | 0.2000 (0.4124) |
| log10(MM) | -0.8808 (0.1954) *** | -0.4363 (0.2933) | -0.8797 (0.2178) *** | -0.3182 (0.3316) |
| log10(LM) | 1.0504 (0.2255) *** | 0.5859 (0.3708) | 1.0737 (0.2606) *** | 0.5317 (0.4290) |
| IP | 0.5916 (0.1124) *** | 0.4381 (0.1754) * | 0.5964 (0.1317) *** | 0.4851 (0.2053) * |
| S | 0.0014 (0.0021) | -0.0004 (0.0019) | 0.0011 (0.0025) | -0.0012 (0.0019) |
| A | 0.0022 (0.0016) | 0.0019 (0.0016) | 0.0024 (0.0023) | 0.0017 (0.0020) |
| Protein (g/100 g) | full $N=35$, tips $R^2=0.812$, tips diet $F$ $P=0.045$, gls diet $F$ $P=0.047$<br>reduced $N=29$, tips $R^2=0.836$, tips diet $F$ $P=0.002$, gls diet $F$ $P=0.007$ | | | |
| (Intercept) | 0.8601 (0.1359) *** | 0.5967 (0.2014) ** | 0.8752 (0.1297) *** | 0.5425 (0.1962) * |
| log10(MM) | -0.3746 (0.1107) ** | -0.2459 (0.1634) | -0.3318 (0.1038) ** | -0.0874 (0.1627) |
| log10(LM) | 0.2476 (0.1285) | 0.1708 (0.2046) | 0.1549 (0.1243) | -0.0676 (0.2093) |
| IP | 0.4157 (0.0636) *** | 0.4494 (0.0985) *** | 0.4528 (0.0628) *** | 0.5143 (0.0952) *** |
| S | 0.0030 (0.0012) * | 0.0025 (0.0010) * | 0.0043 (0.0012) ** | 0.0029 (0.0011) * |
| A | 0.0009 (0.0009) | 0.0012 (0.0009) | 0.0031 (0.0011) ** | 0.0028 (0.0010) * |
| Sugar (g/100 g) | full $N=33$, tips $R^2=0.368$, tips diet $F$ $P=0.072$, gls diet $F$ $P=0.153$<br>reduced $N=28$, tips $R^2=0.519$, tips diet $F$ $P=0.004$, gls diet $F$ $P=0.013$ | | | |
| (Intercept) | 0.9135 (0.0464) *** | 0.9904 (0.0633) *** | 0.8932 (0.0439) *** | 0.9568 (0.0569) *** |
| log10(MM) | 0.0382 (0.0376) | -0.0180 (0.0533) | 0.0239 (0.0354) | -0.0397 (0.0487) |
| log10(LM) | -0.0590 (0.0435) | -0.0137 (0.0657) | -0.0224 (0.0427) | 0.0401 (0.0617) |
| IP | -0.0646 (0.0223) ** | -0.0529 (0.0296) | -0.0718 (0.0213) ** | -0.0650 (0.0268) * |
| S | -0.0007 (0.0004) | -0.0004 (0.0004) | -0.0012 (0.0004) ** | -0.0008 (0.0004) |
| A | -0.0006 (0.0003) | -0.0006 (0.0003) | -0.0013 (0.0004) ** | -0.0011 (0.0004) ** |

\* $P < 0.05$ , \*\* $P < 0.01$ , \*\*\* $P < 0.001$

**Figure 2.**
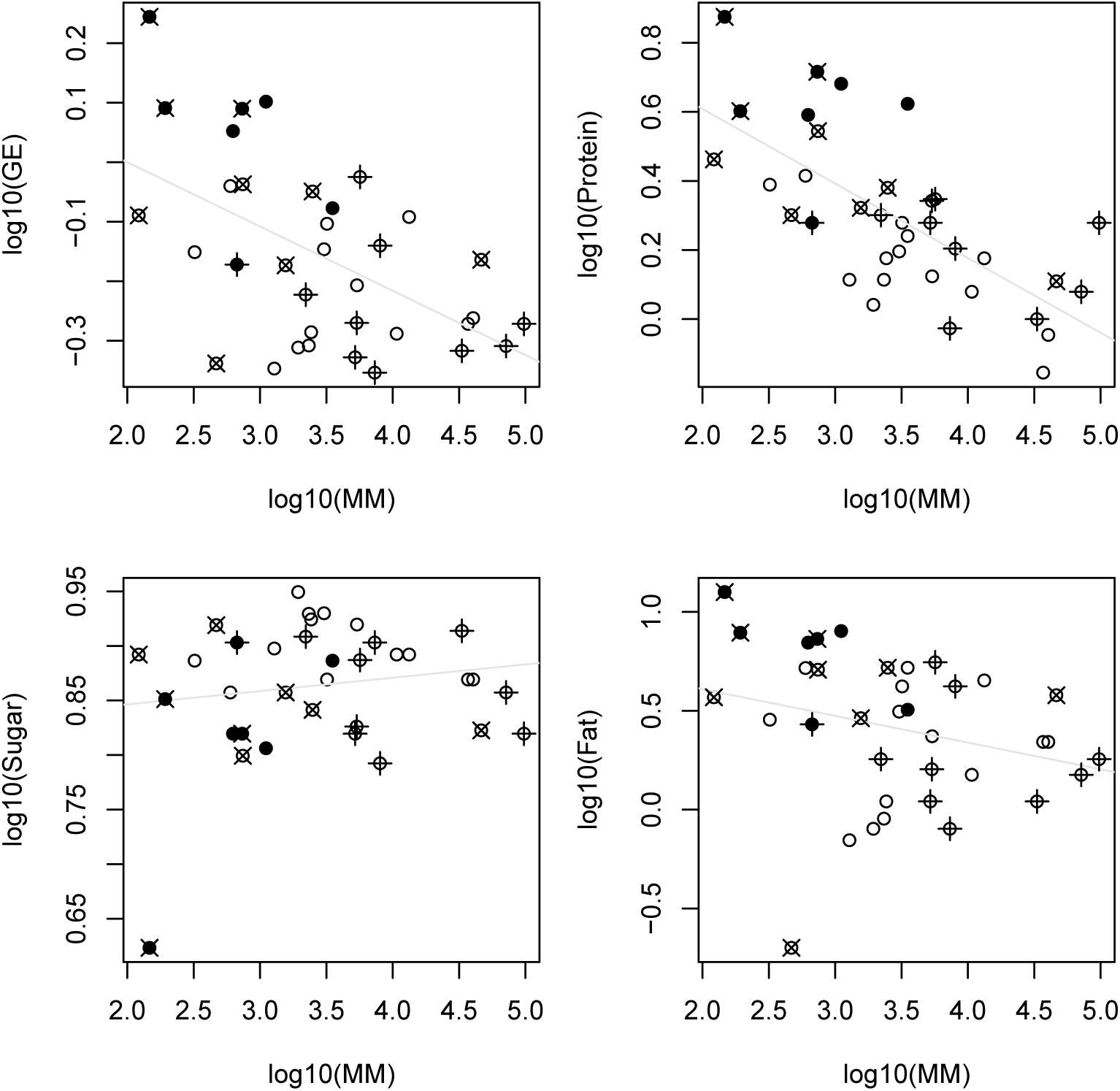
Plots of milk composition against adult body mass. Filled symbols are infant parkers; *×* indicate species with percent animal matter in diet *≥* 30% and + indicate structural plant material *≥* 40%. The extreme low outliers are *Galago moholi* (sugar) and *Callimico goeldii* (fat) were excluded from regression analysis. The only folivorous parker, *Hapalemur griseus*, was also excluded. Regression lines are TIPS bivariate least-squares fits to all the points. TIPS and PGLS results with multiple predictors are given in Table 1.

## Results

Milk gross energy and fat concentration are influenced by the same variables because fat is the primary source of milk energy. In TIPS multiple regressions maternal mass, litter mass, and infant parking are significant predictors of gross energy and fat concentration. Regression slopes indicate decreasing milk energy and fat concentration with increasing maternal size, increasing energy and fat concentration with increasing litter size, and increased energy and fat concentration among infant parkers. However, PGLS models suggest these effects are strongly influenced by phylogeny. Only the infant parking effect remains significant in the PGLS models. Importantly, diet variables are not associated with gross energy or fat concentration in TIPS or PGLS models (Table 1).

In contrast, milk protein and sugar concentration are strongly influenced by diet. Heuristically, the diet effects on milk protein and sugar are seen as displacement of frugivores below (protein) or above (sugar) a TIPS bivariate regression line on maternal mass (Figure 2). The combined-variable diet effect is a significant predictor of protein concentration in the full or reduced (milk samples *>*1) data sets (*F*-tests in Table 1). Protein concentration regression slopes for the individual diet variables are small but significantly positive in all cases, indicating increased structural plant part or animal matter consumption elevates milk protein concentration. This effect persists in PGLS models accounting for phylogeny. Similarly, infant parking significantly elevates protein concentration. Maternal mass is only a significant negative predictor of protein concentration in TIPS multiple regressions.

Milk sugar concentration is not influenced by maternal or litter mass even in TIPS multiple regressions. Infant parking is a significant predictor by TIPS multiple regression in either data set and by PGLS in the reduced data set. The slope is negative indicating that infant parking reduces the sugar concentration of milk. Also, increasing structural plant part or animal matter consumption lowers sugar concentration in the reduced data set (Table 1). Both the diet and parking effects may be an outcome of reduced volume/yield of milk in these less frugivorous and/or less frequently suckling taxa (Hinde & Milligan, 2011).

## Discussion

Our results significantly clarify and extend the known correlates of primate milk composition. We found wild diets to be significant predictors of milk protein and sugar concentration. Higher amounts of structural plant parts or animal matter in the diet increases protein while decreasing sugar concentration. We also found infant parking to be an important predictor of all aspects of proximate milk composition — elevating gross energy, fat concentration, and protein concentration while depressing sugar concentration. This confirms the results of Tilden et al. (Tilden & Oftedal, 1997) but refines them by noting the effect persists even when accounting for phylogenetic relatedness, allometric effects of adult mass, litter mass, and diet. Regression slopes for the diet variables are small. For example, using PGLS slopes for the full data set *β ≈* 0.0025 for percent protein, which implies an increase of about 0.006 to 0.010 percent milk protein for each unit increase in animal or structural plant material in the diet. The infant parking effect is much more dramatic, with a predicted increase of 1.185 percent milk protein for a switch to parking in this model (*β* = 0.45). More precise data on the frequency and duration of inter-nursing bouts caused by the mother-infant separation of parking, or possibly allomaternal carrying (Ross & MacLarnon, 2000; Mitani & Watts, 1997), should clarify our understanding of this relationship. We found little statistical support for strong allometric relationships with maternal size or investment effects via litter mass (Riek, 2008). Maternal mass regression slopes were usually negative, suggesting lower concentrations of protein and fat in larger mothers (Hinde & Milligan, 2011; Martin, 1984). Litter mass slopes were usually positive implying larger neonatal mass requires higher protein and higher fat milk, consistent with mammalian trends of a positive correlation between pre- and postnatal investment (Derrickson, 1992). A larger dataset of primates and other mammals may confirm these to be true biological relationships.

The association between wild diet and milk protein and sugar concentrations could be explained by several overlapping causes including the influence of species-typical foraging patterns, an independent correlation between diet and infant growth rates, variation among foods in the presence of nutrients that are costly to synthesize, and simple transmission of ingested nutrients. The first three of these we consider to be inter-related, plausible explanations which emphasize maternal energetics and can potentially be tested with additional data.

Maternal foraging ecology may explain the milk composition-diet relationship if foods requiring longer foraging time or greater travel distance (e.g. small, patchily distributed fruits) limit the energy mothers can allocate to milk production. A difficulty for this hypothesis is comparative studies of primate travel distance suggest energetic expenditures on locomotion are a small portion (< 5%) of daily energy needs (Steudel, 2000; Chapman & Chapman, 2000). However, energetic costs of locomotion may be greatly underestimated (Altmann, 1987) and intrapopulation variation even around these small values may still have consistent fitness effects, particularly for lactating mothers who are likely to face higher energetic costs of locomotion when encumbered by dependent infants (Altmann & Samuels, 1992). We consider this explanation plausible and potentially testable with inter-or intraspecific data on travel distance, activity budgets, food selection, and maternal mass depletion. Primate mothers appear to employ multiple strategies to meet the costs of lactation by increasing gross energy intake, reducing expenditures, or temporarily relying on stored reserves (cercopithecoids: Altmann & Samuels, 1992; Altmann, 1983; Dunbar & Dunbar, 1988; Koenig et al., 1997; Barrett et al., 2006), (hominoids: Murray et al., 2009; Bates & Byrne, 2009; Pontzer & Wrangham, 2006; Lappan, 2009), (platyrrhines: Guedes et al., 2008; Rose, 1994; Boinski, 1988; Miller et al., 2006; Nievergelt & Martin, 1999; Tardif, 1994), (strepsirhines: Saito, 1988; Vasey, 2005; Sauther, 1994).

A closely related explanation is the possibility that milk composition covaries with primate diets because of (semi-)independent dietary effects on infant growth patterns. That milk properties should be related to growth patterns is unremarkable, but the reasons infant growth and diet covary may be different, or at least of differing strengths, from those that determine the milk-diet relationship. Among these are extrinsic mortality risks including infanticide, limitations of brain growth rates, and aspects of infant/juvenile foraging and nutritional ecology including relative acceleration/delay of digestive system maturation and availability of foods for complementary feeding (Lee, 1999; Martin, 1996; Janson & van Schaik, 1993; Leigh, 2004; Charnov & Berrigan, 1993; Godfrey et al., 2001; Langer, 2008, 2003). Differentiating this growth hypothesis from maternal energetics may amount to distinguishing between direct and indirect selection on mothers and their growing offspring (Arnold, 1994). Has selection operated directly on maternal energetics, offspring growth rates, or both and what are the indirect consequences of selection on one of these on the other (Cheverud & Wolf, 2009)? Our results provide general support for Leigh’s (1994) prediction on the high protein content of folivorous primate milk. However, because of this complicated network of possible causes of the milk composition – diet relationship and the paucity of folivores in our data set, we feel it is premature to assess its underlying mechanism or generality. Interestingly, *Callimico* and *Hapalemur*, both small-bodied primates with diets high in structural plant parts, were two of the most problematic points in the dataset, which points toward possible heterogeneity in milk properties among primate folivores (Leigh, 1994).

Finally, it is possible that maternal dietary intake may have a more direct influence the composition of milk by supplying specific nutrients that are transmitted to offspring or by affecting the energetics of milk synthesis. Consistent with our results, structural plant and animal foods are typically considered higher in protein than most fruits. Similarly, fruits are often considered rich sources of sugars and water but little else. However, the lack of a relationship between animal sources and fat content is inconsistent with this explanation and highlights general problems with this hypothesis. A strict interpretation of the relationship between consumed food and milk composition is complicated by the fact that milk is a synthesized product rather than a passive transfer mechanism for nutrients in the form they are encountered in foods. Body reserves are mobilized to varying degrees by mammalian mothers to fuel lactation (Dufour & Sauther, 2002; Gittleman & Thompson, 1988; Tardif et al., 2001; Jönsson, 1997; Stephens et al., 2009). Fatty acids in milk are something of an exception in that they do often directly reflect consumed foods (Milligan & Bazinet, 2008; Milligan et al., 2008). Nevertheless, we found no interspecific relationship between wild diets and milk total fat concentration.

A less strict interpretation of the relationship between consumed foods and milk production, which is more plausible, is that particular nutrients are costly to synthesize and consuming them in larger quantities enables mothers to elevate their concentrations in milk by altering her energy budget (Leigh, 1994; Power et al., 2002; Kirkwood, 1985). A current difficulty of this interpretation, is that the dietary categories of structural and reproductive plant parts, and percents time feeding on them, are often poor indicators of the nutritional composition of ingested foods (Lambert, 2011; Ganzhorn et al., 2009; Chapman et al., 2003; Oftedal, 1991). However, this more general interpretation is compatible with the maternal foraging ecology hypothesis described above, but it emphasizes the benefits of high concentrations of limiting nutrients in foods rather than the energetic costs of finding and processing them while encumbered with an infant. More precise data on the nutrient composition of ingested foods would help define the scope of these potentially complementary hypotheses.

Using milk samples from captive individuals is potentially problematic, but we found no systematically confounding effects of captivity on milk composition. Captivity is likely to increase the average energy available to mothers through rich diets and lowered activity levels, and reduce variation by provisioning of a regular food supply. Within our sample, a handful of available comparisons suggest very limited effects of captivity on the diet-related trends identified in our analysis. The only true folivores (*Gorilla beringei* and *Alouatta* sp.) in the analysis were sampled in wild populations. Howler monkey values for protein are lower than expected and inconsistent with the general dietary pattern, perhaps reflecting reduction due to wild environment. Without comparable data collected in captivity it is unclear if this is the case or not. The contrast between gorilla taxa is consistent with the dietary trend identified, but implies no effect of captivity as it is the wild animals producing richer milk. Finally, at small size where captivity effects are expected to be largest (Power et al., 2008), if only wild values were used for the primarily frugivorous *Callithrix* and *Leontopithecus*, assuming this better reflects their evolved milk properties, it will strengthen the identified milk-diet relationships by reducing protein and raising sugar concentration.

In conclusion, we see no reason to doubt that wild primate diets predict the protein and sugar concentrations of their milk. While the mechanisms linking diet and milk composition are currently unclear, simple transmission of nutrients from consumed foods into milk is the least likely. Instead, three stronger candidate explanations focus on details of maternal energetics. In addition to a larger set of species with milk and life history data used in this analysis, more precise data on food consumption and the nutritional properties of foods, documenting ontogenetic variation among species in infant mass or key organ development, and the continued study of the behavioral ecology of motherhood will resolve the relative effects on primate milk composition of costly nutrients, infant growth, maternal foraging ecology.

## Acknowledgements

We thank Chris Whitter for sharing mountain gorilla neonate mass data with us. Steve Leigh and Charlie Nunn provided helpful comments on the manuscript.

## Supporting Information

Supplementary tables and data source references. (pdf)

Spreadsheet of data used in the analysis. (xls)

Nexus formatted phylogeny used in the analysis. (txt)

